# A TCER-1-siRNA Regulatory Axis Suppresses Antibacterial Innate Immunity in *C. elegans*

**DOI:** 10.64898/2026.01.31.702960

**Authors:** Nikki Naim, Francis RG Amrit, Laura L Bahr, Brooke E. Montgomery, Spencer Kuhn, Taiowa A. Montgomery, Arjumand Ghazi

## Abstract

Small interfering RNA (siRNA) are important regulators of gene expression with well-established roles in pathogen defense. Yet, their specific roles in antibacterial immunity are not well understood. Here, we identify an siRNA pathway involved in repressing antibacterial innate immunity in *Caenorhabditis elegans*. We show that genes required for the biogenesis or function of WAGO Argonaute-associated siRNAs, called 22G-RNAs, function in a common genetic pathway with the immune-suppressive transcription elongation and splicing factor TCER-1 to inhibit immunity. Loss of *tcer-1* reduced levels of 22G-RNAs from a subset of WAGO targets, while mutations in several WAGO 22G-RNA pathway genes phenocopied the enhanced immunoresistance of *tcer-1* mutants, suggesting a shared regulatory module. Integrative 22G-RNA-mRNA analyses and molecular genetic studied show that this module does not induce widespread gene silencing, but instead targets a restricted set of immune-relevant effectors, including *scrm-4*, encoding a phospholipid translocase that promotes host resistance. Together, our findings establish endogenous WAGO 22G-RNAs as repressors of antibacterial immunity and identify TCER-1 as a physiological regulator that promotes 22G-RNA biogenesis to constrain host defense. The results uncover a previously unrecognized small RNA-dependent mechanism linking transcription, metabolism, and antibacterial innate immunity.

## INTRODUCTION

Non-coding small RNAs, including microRNAs (miRNAs) and siRNAs, mediate RNA interference (RNAi)-based gene silencing to regulate diverse processes ranging from development to genome defense [1, 2]. miRNAs are widely established as conserved regulators of innate and adaptive immune responses against viral and bacterial pathogens, whereas siRNAs have roles in antiviral immunity [3–7]. In plants and invertebrates, Dicer-mediated processing of viral double-stranded RNA and subsequent Argonaute-dependent silencing constitute a canonical, genetically well-defined antiviral pathway [7–10]. In mammals, miRNAs have well-defined roles in shaping innate and adaptive immune programs [3, 4, 6], whereas functions for endogenous siRNAs remain poorly defined [11–14]. Similarly, although small RNAs contribute to antibacterial responses in plants and invertebrates, these functions have largely been described in the context of miRNAs [15–18].

The nematode *Caenorhabditis elegans*, despite lacking professional immune cells, deploys defense programs regulated by conserved signaling cascades[19]. miRNA-based antibacterial defenses have been described in *C. elegans*, along with roles for specific Argonaute proteins [18, 20–25]. In *C. elegans*, endogenous siRNAs are dominated by two branches defined by their Argonaute interactors: CSR-1 and the WAGOs, which together comprise several distinct proteins [2]. The CSR-1 22G-RNA branch licenses germline gene expression, whereas the WAGO 22G-RNA branch mediates exogenous and endogenous RNAi, including repression of aberrant transcripts and transposons [26]. Previous work has revealed roles for the piwi-interacting RNA (piRNA) and downstream RNAi pathway in bacterial sensing and pathogen-avoidance learning [27–29]. Recently, CSR-1 was shown to promote antibacterial resistance [20], but how distinct Argonaute-associated small RNA pathways contribute to antibacterial immunity remains poorly defined.

We previously identified TCER-1, the *C. elegans* homolog of human transcription elongation and splicing factor TCERG1 [30–34], as a repressor of host defense against multiple Gram-positive and Gram-negative pathogens, as well as abiotic stressors [35]. *tcer-1* mutants exhibit enhanced immunoresistance, whereas its overexpression shortens post-infection survival [35]. Mutants of the *Arabidopsis thaliana* TCER-1 homolog, PRP40, also show increased pathogen resistance[36]. TCER-1 has emerged in genetic screens for RNAi regulators suggesting that it may suppress immunity via a small RNA pathway [37, 38]. Here, we show that *tcer-1* mutants exhibited reduced levels of 22G-RNAs from a subset of WAGO targets, and mutations in several WAGO genes phenocopied the enhanced immunoresistance of *tcer-1* mutants against the human opportunistic pathogen *P. aeruginosa* PA14. Integrative small RNA (sRNA)-mRNA sequencing revealed that this pathway does not induce global gene silencing but targets select immune-relevant effectors, including *scrm-4*, encoding a phospholipid translocase, that promotes host defense. Together, our findings establish endogenous WAGO 22G-RNAs as repressors of antibacterial immunity and identify TCER-1 as a physiological regulator that promotes 22G-RNA biogenesis to constrain host defense.

## RESULTS

### Overlapping roles for TCER-1, PPW-1, and RRF-1 in suppressing anti-bacterial immunity

TCER-1 is broadly expressed in the soma and germline [35, 39]. Thus, to identify the tissues in which TCER-1 is necessary to repress immunity, we used *ppw-1(pk1425)* and *rrf-1(ok589)* mutants, which restrict RNAi to the soma and germline, respectively. *ppw-1* encodes an Argonaute and *rrf-1* encodes an RNA-directed RNA polymerase; both genes function specifically within the WAGO 22G-RNA pathway and strains containing mutations in these genes are widely used for tissue-restricted RNAi in worms [40, 41]. Unexpectedly, both *ppw-1* and *rrf-1* mutants survived significantly longer upon PA14 infection compared to wildtype animals (**Fig. 1A-B, Supplementary Table S1**). Moreover, *tcer-1* RNAi did not further enhance post-infection survival in either mutant (**Fig. 1A-B, Supplementary Table S1**). Additionally, mutating *ppw-1* or *rrf-1* did not significantly further extend PA14 survival in *tcer-1(tm1452)* mutants, suggesting that PPW-1 and RRF-1 may regulate immunity through the same pathway as TCER-1 (**Fig. 1C-D, Supplementary Table S2**). Given their roles in gene regulation, we tested whether *tcer-1*, *rrf-1*, or *ppw-1* regulate each other’s expression as a potential explanation for their shared role in PA14 resistance. GFP expression from a TCER-1::GFP translational reporter [35, 39] was unchanged in either the *ppw-1* or *rrf-1* mutant (**Fig. 1E**). Conversely, *ppw-1* mRNA levels were unchanged in *tcer-1* mutants, whereas *rrf-1* mRNA levels were modestly reduced (∼1.5-fold) based on the RNA-seq analysis described below, which may indicate that TCER-1 acts upstream of *rrf-1* to suppress antibacterial resistance.

**Figure 1.**
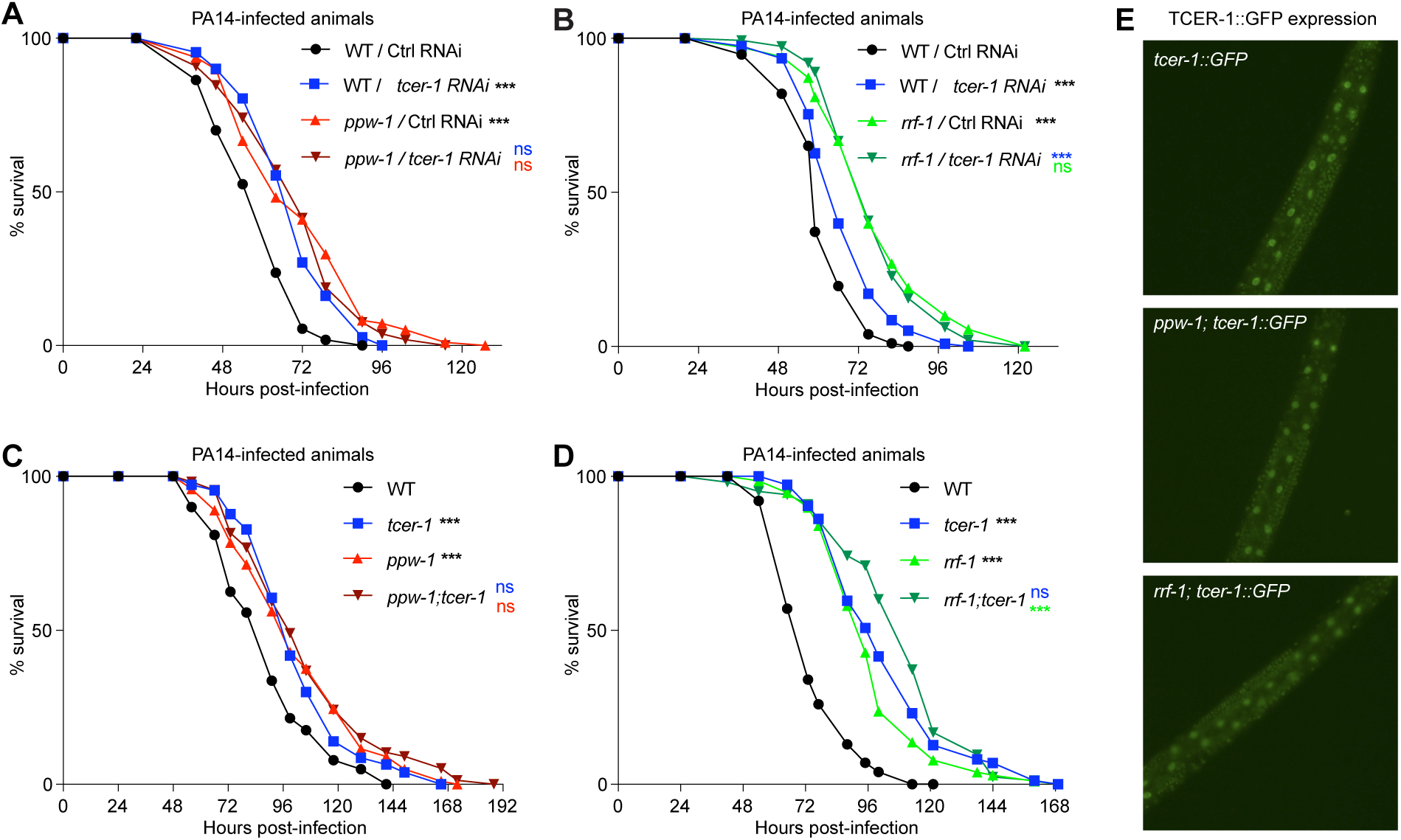
TCER-1, RRF-1, and PPW-1 act nonadditively to regulate antibacterial immunity. **(A-D)** Impact of *tcer-1* inactivation by RNAi (A, B) or mutation (C, D) on survival of wildtype (WT) animals and *ppw-1* or *rrf-1* mutants following PA14 exposure. **A, B:** WT, *ppw-1(pk1425)* and *rrf-1(ok589)* grown from egg to L4 larval stage on bacteria expressing dsRNA targeting the control empty vector (Ctrl) or *tcer-1* and exposed to PA14 from L4 onwards. **A:** WT, Ctrl (black: m= 58.52 ± 1.48, n= 63/81); WT, *tcer-1* RNAi (blue, m= 69.23 ± 1.48, n= 78/109); *ppw-1* Ctrl RNAi (red, m= 71.2 ± 1.95, n= 99/112), *ppw-1*, *tcer-1* RNAi (maroon, m= 70.14 ± 1.62, n= 110/121). **B:** WT, Ctrl RNAi (black, m= 60.93 ± 0.95, n= 110/134); WT, *tcer-1* RNAi (blue, m= 67.91 ± 1.09, n= 129/157); *rrf-1*, Ctrl RNAi (neon green, m= 77.45 ± 1.67, n= 114/136); *rrf-1*, *tcer-1* RNAi (olive green, m= 77.3 ± 1.33, n= 117/154). **C-D:** Survival of WT and mutant strains upon PA14 exposure from L4 stage onwards. **C:** WT (black, m= 88.97 ± 2.01, n= 116/143), *tcer-1(tm1452)* (blue, m= 102.76 ± 2.3, n= 96/111), *ppw-1(pk1425)* (red, m= 102.95 ± 2.38, n= 134/148), *ppw-1(pk1425);tcer-1(tm1452)* (maroon, m= 106.68 ± 2.94, n= 95/116). **D:** WT (black, m= 74.9 ± 1.34, n= 108/130), *tcer-1(tm1452)* (blue, m= 103.91 ± 2.45, n= 91/120), *rrf-1(ok589)* (neon green, m= 97.99 ± 1.89, n= 113/137), *rrf-1(ok589);tcer-1(tm1452)* (olive green, m= 108.5 ± 2.55, n= 88/106). **E:** Representative images of TCER-1::GFP expression in WT animals and *ppw-1(pk1425)* and *rrf-1(ok589)* mutant strains. In A-D, mean survival following PA14 exposure shown in hours (m) ± SEM; n= observed/total (see Methods for details). Significance was calculated and using the log-rank method (Mantel Cox, OASIS2), and *p* values were adjusted for multiple testing using the Bonferroni procedure. Statistical significance shown on each panel with the color of the asterisk indicating the strain being compared to. *p* <0.05 (*), < 0.01 (**), <0.001 (***), not significant (ns). Data from additional trials are presented in Supplementary Table S1.

### Multiple WAGO 22G-RNA biogenesis factors suppress immunoresistance

To further explore the role of 22G-RNAs in the PA14 response, we examined PA14 resistance in several additional WAGO pathway mutants - *mut-16*, *mut-14* and *smut-1* (which function partially redundantly), *mut-7*, and *rde-3* - all of which encode core components of the Mutator complex (**Fig. 2A**) [42–47]). With the exception of *rde-3*, all exhibited enhanced PA14 resistance comparable to *tcer-1* mutants, although *mut-16* showed a more modest effect across trials (**Fig. 2B-E, Supplementary Table S2**). RDE-3 is a nucleotidyltransferase that promotes entry of cleaved mRNAs into the 22G-RNA pathway via poly(UG) tailing [48]. The lack of enhanced survival in *rde-3* mutants suggests that this activity may not be for required suppressing antibacterial resistance, or that it also performs functions that offset this effect. In contrast to WAGO 22G-RNA factors, mutations in *rde-1*, which encodes an Argonaute required for exogenous 22G-RNA production and antiviral RNAi, but which is largely dispensable for endogenous 22G-RNAs [49–56], reduced PA14 survival relative to wildtype (**Fig. 2F, Supplementary Table S2**). This indicates that the WAGO 22G-RNA pathway’s role in regulating antibacterial resistance is distinct from its role in the antiviral RNAi pathway involving RDE-1.

**Figure 2.**
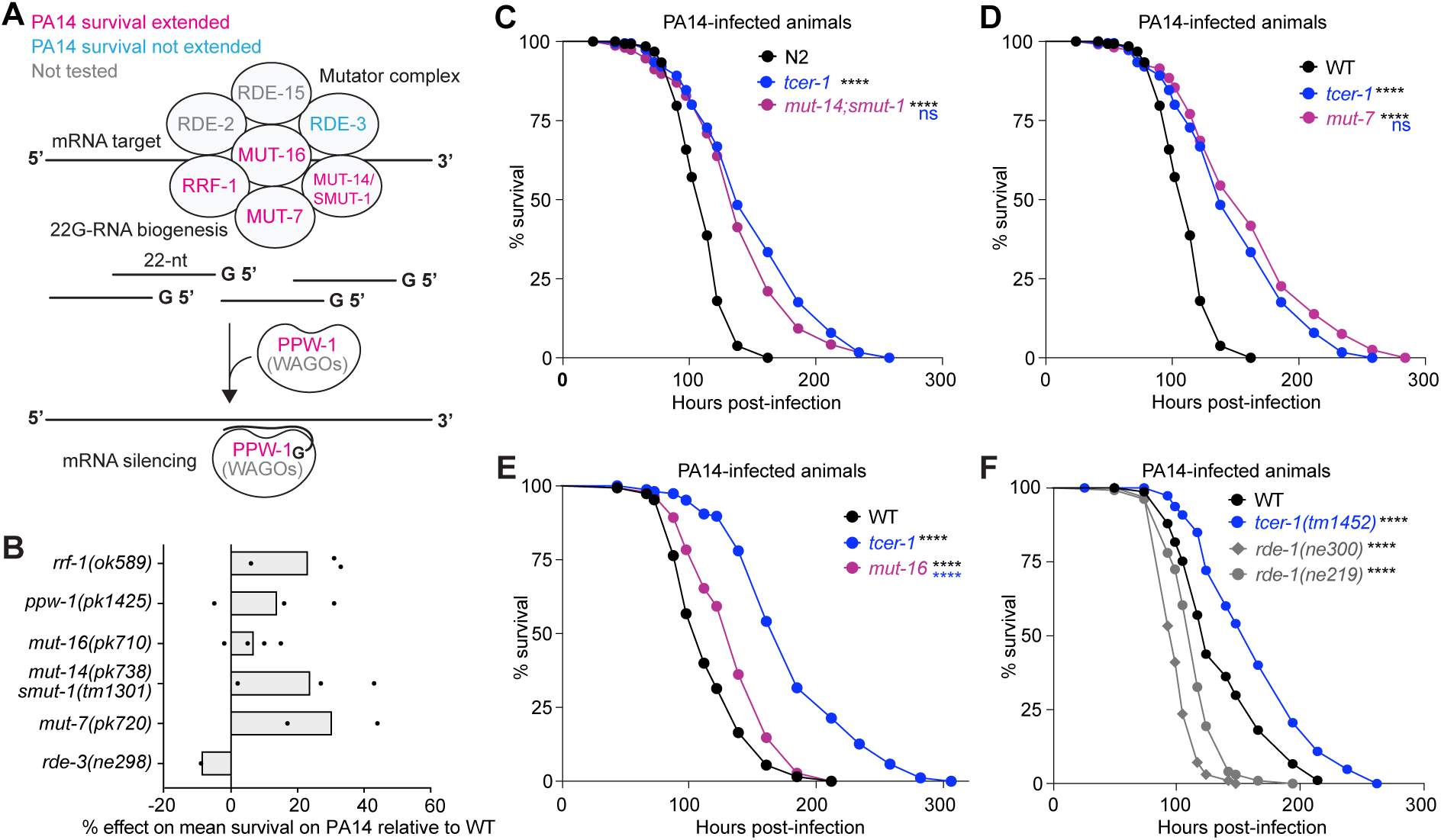
Loss of WAGO 22G-RNA biogenesis genes leads to enhanced immunoresistance. **A:** Schematic of the WAGO 22G-RNA pathway. **B:** Percent effect of 22G-RNA biogenesis factor mutations on survival upon PA14 infection compared to WT control (black line). Each data points represents percent effect in an independent trial. **C-F:** Representative graphs showing survival of *mut-14 smut-1* (C), *mut-7* (D), *mut-16* (E) and *rde-1* (F) mutants following PA14 infection compared to WT animals and *tcer-1* mutants. **C:** WT (m= 111.5 ± 1.9, n= 112/162), *tcer-1(tm1452)* (m= 151.4 ± 4.2, n= 120/201), *mut-14(pk738) smut-1(tm1301)* (m= 141.8 ± 3.7, n= 134/160). **D:** WT (m= 111.5 ± 1.9, n= 112/162), *tcer-1(tm1452)* (m= 151.4 ± 4.2, n= 120/201), *mut-7(pk720)* (m= 160.9 ± 5.5, n= 89/115). **E:** WT (m= 118.2 ± 2.7, n= 137/155), *tcer-1(tm1452)* (m= 183.11 ± 4.3, n= 119/170), *mut-16(pk710)* (m= 135.95±2.9, n= 127/145). **F:** WT (m= 137.64 ± 3.3, n= 117/158), *tcer-1(tm1452)* (m= 166.53 ± 4.4, n= 94/167), *rde-1(ne300)* (m= 102.26 ± 1.2, n= 105/134), *rde-1(ne219)* (m= 115.25 ± 2.0, n= 105/135). In C-F, survival following PA14 exposure is shown in mean hours (m) ± SEM; n= observed/total (see Methods for details). Significance was calculated and using the log-rank method (Mantel Cox) and *P* values were adjusted for multiple testing using the Bonferroni procedure. Statistical significance shown on each panel with the color of the asterisk indicating the strain being compared to. *p* <0.0001 (***), not significant (ns). Data from additional trials and other details are in Supplementary Table S3.

### *tcer-1* is required for normal 22G-RNA accumulation at a subset of WAGO targets

Based on our observations above and prior reports implicating TCER-1 in the WAGO 22G-RNA pathway [37, 38], we asked whether TCER-1 is required for the formation or accumulation of WAGO 22G-RNAs and whether it also plays a role in regulating WAGO target mRNAs. To address these questions, we performed high-throughput sequencing of small RNAs and mRNA from age-matched day 1 adults of *tcer-1(tm1452)* mutants and wildtype animals grown on non-pathogenic *E. coli* OP50 (**Fig. 3A**). Because *tcer-1(tm1452)* mutants exhibit developmental asynchrony that can potentially confound RNA-seq analysis due to stage and tissue specific gene expression [57], we included an independently generated CRISPR deleted allele, *tcer-1(glm27)*, which phenocopies *tcer-1(tm1452)*-including enhanced PA14 resistance-but lacks developmental asynchrony (**Fig. S1**).

**Figure 3.**
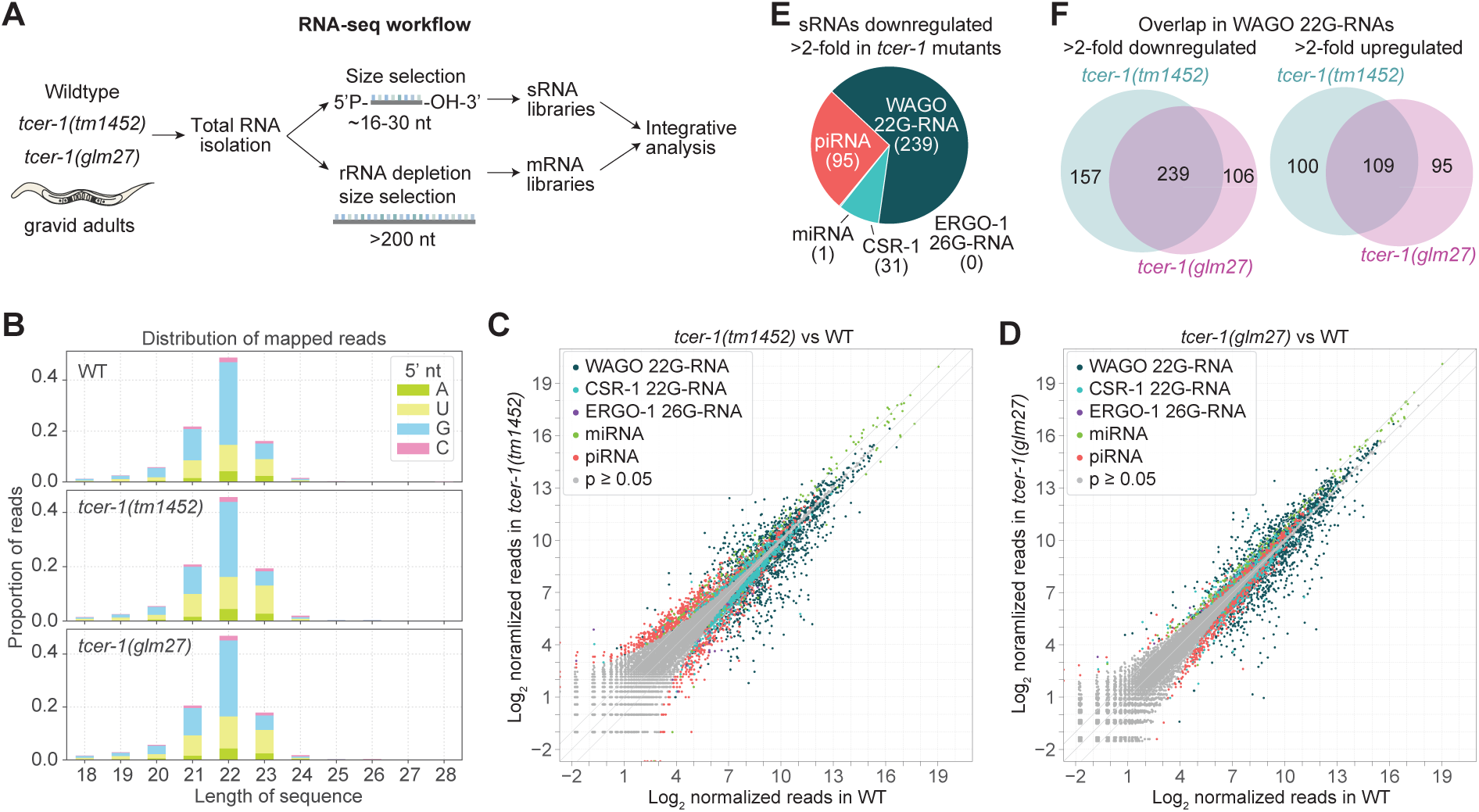
*tcer-1* inactivation leads to depletion of a subset of WAGO 22G-RNAs. **A:** Schematic of tandem sRNA-mRNA-seq pipeline. **B:** Size and 5’-nt distribution of sRNA-seq reads in WT and *tcer-1* mutants. One of 3 biological replicates is shown for each strain. **C-D:** Scatterplots showing individual small RNA-producing genes as the average log_2_ geometric mean (GM)-normalized sRNA-seq reads in WT animals (x axis) and *tcer-1(tm1452)* (C) or *tcer-1(glm27)* mutants (D) (y axis). *p* values were calculated using the Wald test with DESeq2. Small RNAs are color-coded by class. **E:** Pie chart showing the classification of the 366 small RNA features reduced in both *tcer-1* mutants. **F:** Overlap of WAGO targets showing >2-fold enrichment or depletion of 22G-RNAs in *tcer-1* mutants. See Supplementary Table S4 for differential expression analysis of sRNAs.

Global small RNA profiles were highly similar between wildtype and *tcer-1* mutants, and were dominated by 22-nucleotide (nt), 5’G-containing sequencing reads characteristic of 22G-RNAs [51], indicating that *tcer-1* does not broadly dictate the small RNA landscape (**Fig. 3B**). Small RNAs were classified as miRNAs, piwi-interacting RNAs (piRNAs), or siRNAs, with siRNAs subclassified by their associated Argonautes (CSR-1, WAGO, or ERGO-1) and grouped by the gene from which they were processed [26]. Both *tcer-1* alleles produced highly concordant effects on small RNA abundance **(Fig. 3C-D and Supplementary Table S4).** A total of 366 small RNA features were reduced in both mutants, 270 (74%) of which were 22G-RNAs. Among these 270 affected 22G-RNA loci, 31 (11%) corresponded to CSR-1 targets, whereas 239 (89%) were WAGO targets (89%) (**Fig. 3E, Supplementary Table S4**). Of the 2,215 annotated WAGO targets analyzed [43], 348 were misexpressed in *tcer-1* mutants, including the 239 that were downregulated and an additional 109 that were upregulated in both *tcer-1* alleles (**Fig. 3F, Supplementary Table S4**). These data indicate that TCER-1 selectively influences 22G-RNA formation or stability from a subset of WAGO targets. Although these effects may be indirect, possibly caused by altered processing of WAGO target mRNAs in *tcer-1* mutants, our genetic epistasis analyses are consistent with a model in which TCER-1 and WAGO 22G-RNAs function through a shared set of targets to suppress immunity.

### TCER-1 functions through the 22G-RNA target *scrm-4* to repress antibacterial resistance

Using the mRNA-seq data generated in parallel with the sRNA-seq data, we assessed the impact of *tcer-1* loss on mRNA expression of WAGO 22G-RNA target genes [43, 44]. None of the known genes involved in WAGO 22G-RNA biogenesis or amplification were significantly downregulated >2-fold in both *tcer-1* mutants, indicating that misexpression of WAGO pathway genes is unlikely to underlie the observed loss of 22G-RNAs. However, more modest changes, such as the ∼1.3-fold downregulation of *rrf-1*, could contribute to the phenotype (**Supplementary Table S5**). Both *tcer-1* mutants exhibited modest, bidirectional changes in WAGO target gene expression; of the 2,215 annotated WAGO targets only a small fraction were consistently misexpressed in both mutants, with 90 upregulated and 34 downregulated (**Fig. 4A-C, Supplementary Table S5**).

**Figure 4.**
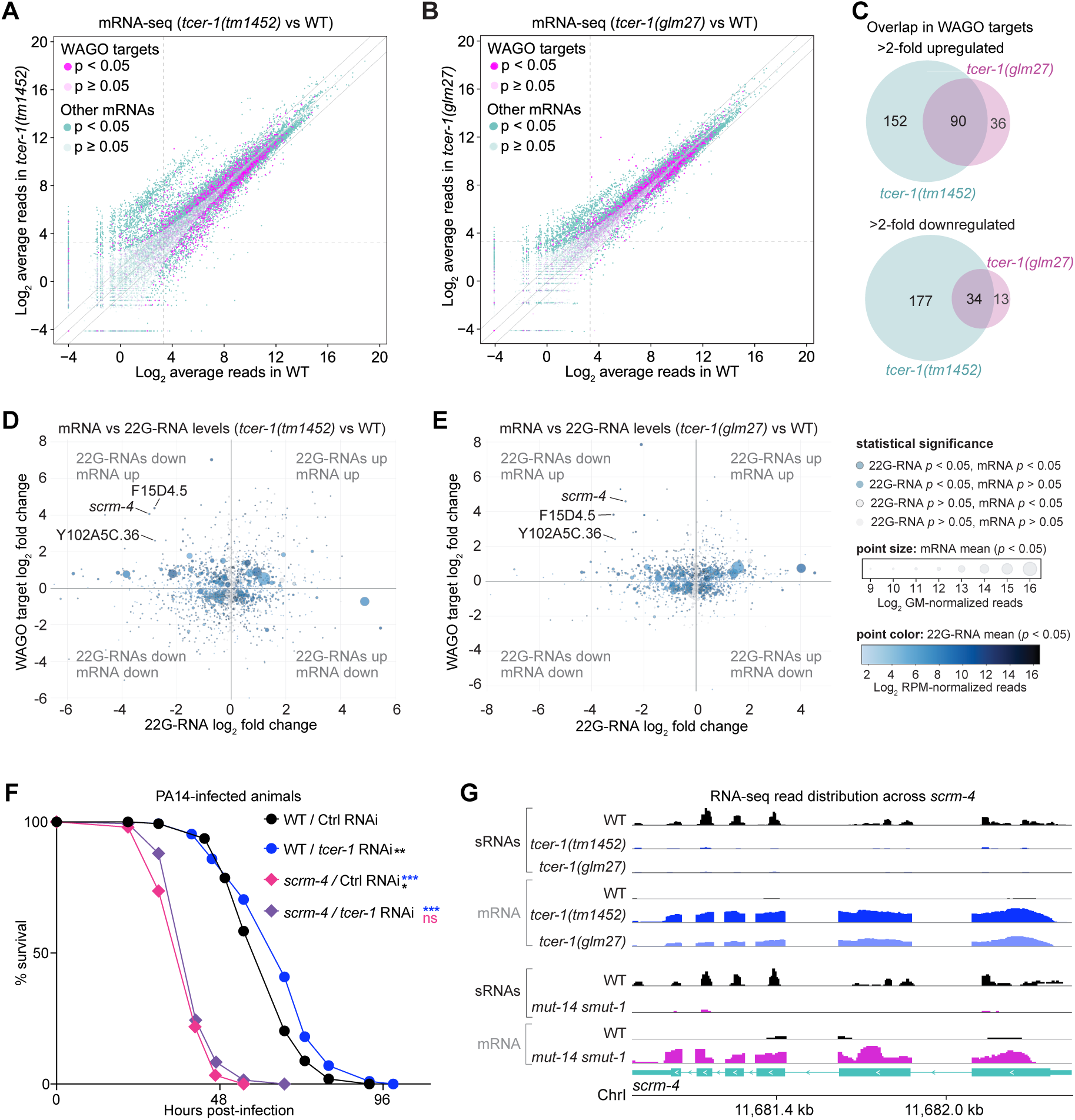
Mutation of the TCER-1 and WAGO 22G-RNA target *scrm-4* compromises antibacterial immunity. **A-B:** Scatterplots showing WAGO 22G-RNA producing genes as the average log_2_ GM-normalized mRNA-seq reads in wildtype (WT, x axis) and *tcer-1(tm1452)* (A) or *tcer-1(glm27)* mutants (B) (y axis). n=3 biological replicates. **C:** Overlap in WAGO 22G-RNA producing genes enriched or depleted of mRNA-seq reads >2-fold in *tcer-1* mutants. See Supplementary Table S5 for differential expression analysis of all mRNA features. **D-E:** Correlation between WAGO 22G-RNAs and target mRNAs. Cosmic plots displaying WAGO 22G-RNA producing genes as a function of log_2_ fold change in mRNA levels (y axis) and 22G-RNA levels (x axis) in *tcer-1(tm1452)* (D) or *tcer-1(glm27)* mutants (E). Read abundance and statistical significance is indicated as outlined in the key in (E). **F:** Survival of WT and *scrm-4(ok3596)* mutants grown from egg to L4 larval stage on empty vector control (Ctrl) or *tcer-1* RNAi bacteria and subsequently exposed to PA14. WT/Ctrl RNAi (mean= 62.42 ± 1.0, n= 118/150), WT/*tcer-1* RNAi (mean 66.42 ± 1.22, n= 112/157), *scrm-4*/Ctrl RNAi (pink, m= 39.29 ± 0.57, n= 141/149), *scrm-4*/*tcer-1* RNAi (purple, m= 41.86 ± 0.56, n= 144/150).

Given the disparity between the numbers of misregulated 22G-RNAs and WAGO target mRNAs in *tcer-1* mutants, we next assessed the relationship between WAGO target mRNA levels and corresponding 22G-RNA abundance. There was no clear global anticorrelation between the mRNAs and 22G-RNA levels, consistent with similarly weak anticorrelation in *mut-16*, *wago-1*, and *wago-3* mutants in prior reports (**Fig. 4D-E, Supplementary Tables S6-S7)**[58, 59]. Only 14 WAGO targets exhibited the expected anticorrelative relationship between 22G-RNA and mRNA abundance and had both 22G-RNAs and mRNAs that were misexpressed >2-fold in both *tcer-1* mutants (**Supplementary Tables S6-S7**). Among these, only three genes, F15D4.5, Y102A5C.36, *scrm-4*, showed particularly strong effects, with 22G-RNAs reduced >4-fold and corresponding mRNA levels increased by >4-fold (**Fig. 4D-E**, **Supplementary Tables S6-S7**). F15D4.5 and Y102A5C.36 lack conserved protein domains, whereas *scrm-4* encodes a phospholipid scramblase orthologous to human PLSCR3, which regulates membrane phospholipid translocation, mitochondrial structure and function, and plays a role in apoptosis [60–62]. Given the established role of lipid metabolism in *C. elegans* immunity [63] and our recent work demonstrating TCER-1-mediated phospholipid remodeling in immunometabolism [64], we tested whether *scrm-4* contributed to the PA14 response. *scrm-4* loss-of-function mutants survived significantly shorter that wildtype upon PA14 exposure (**Fig. 4F**). Moreover, *tcer-1* RNAi failed to extend *scrm-4* mutants’ survival on PA14 suggesting that *tcer-1* functions through *scrm-4* to control PA14 response (**Fig. 4F**). Notably, changes in *scrm-4* mRNA and 22G-RNA levels were similar in *tcer-1* and *mut-14 smut-1* mutants, suggesting a shared role for TCER-1 and the WAGO 22G-RNA pathway in regulating *scrm-4* expression (**Fig. 4G**). Together these data suggest that TCER-1 and the WAGO 22G-RNA pathway downregulate SCRM-4 to suppress antibacterial immunity.

## DISCUSSION

In this study, we identified WAGO 22G-RNAs as suppressors of antibacterial immunity in *C. elegans*. Our results suggest that PPW-1 is the WAGO Argonaute that modulates this response, consistent with a recent study that also showed *ppw-1* mutants have increased PA14 resistance [23]. Given the functional redundancy amongst WAGO Argonautes, it is possible that additional WAGOs function in this pathway as well[23, 51, 65]. We showed that the WAGO 22G-RNA pathway acts within an immune-suppressive regulatory program involving the transcription elongation and splicing factor TCER-1. Furthermore, our integrated sRNA- and mRNA-sequencing, together with *in vivo* molecular genetic analyses suggest that TCER-1 and the WAGO pathway regulate immunity through selective targeting of specific, immunologically relevant genes rather than through broad siRNA-mediated silencing. One such target is *scrm-4*, a lipid-metabolic gene that contributes to host defense in a TCER-1-dependent manner.

The piRNA pathway promotes *P. aeruginosa* avoidance in *C. elegans*, thereby protecting animals from pathogen infection [27–29]. In contrast, our results indicate that the WAGO 22G-RNA pathway has the opposite effect on pathogen susceptibility by suppressing immunoresistance. These two pathways are likely distinct, since the small RNAs reduced in *tcer-1* mutants are predominantly WAGO 22G-RNAs, whereas those depleted following PA14 exposure are largely piRNAs. Furthermore, loss of the piRNA biogenesis factor *prde-1* did not enhance survival on PA14 (Supplementary Table S3). Moreover, our prior work showed that *tcer-1* mutants’ immunoresistance is not due to reduced pathogen intake [35], supporting a model in which TCER-1 regulates post-infection molecular immunity.

Small RNA-based regulation of antibacterial immunity is an emerging concept across species [13, 17, 23, 66]. Although WAGO Argonautes are not conserved outside of nematodes, components of this pathway, such as *mut-7*, are conserved from worms to humans [42], raising the possibility that analogous regulatory mechanisms-potentially involving TCER-1 orthologs-link small RNA pathways to antibacterial immunity in other species as well.

## METHODS

### *C. elegans* strains and culture

Worm strains were maintained on NGM seeded with *E. coli* strain OP50 using standard methods [67]. RNAi experiments were conducted on NGM supplemented per liter with 1 ml of 1 M IPTG and 1 ml of 100 mg/ml ampicillin and seeded with *E. coli* strain HT115 expressing RNAi constructs or empty vector (pAD12). For FUDR experiments, NGM was supplemented with 100 μg/ml FUDR [35].

### Pathogen stress assays

Pathogen survival assays were performed using the *P. aeruginosa* PA14 slow-killing model [35, 69, 70]. PA14 was streaked from -80°C stocks onto LB agar, incubated overnight at 37°C, stored at 4°C (≤1 week), and single colonies were grown in King’s Broth (16–18 h, 37°C, shaking). Cultures (20 µL) were seeded onto slow-killing plates (modified NGM, 0.35% peptone), incubated at 37 °C for 24 h and then at room temperature for 24 h. Age-matched animals were grown on OP50 or HT115-RNAi plates to L4, designated Day 0 of adulthood, and transferred (≥100 animals; 20–30/plate) to five PA14 plates per strain. For FUDR assays, L4 animals were placed on FUDR plates for 24 h at 15°C prior to transfer. Survival was scored every 6–12 h at 25°C or 20°C, with daily transfers to fresh PA14 plates for the first 3–4 days to separate adults from progeny. Animals that exploded, bagged, crawled off plates, or became contaminated were censored. Kaplan–Meier survival analyses were performed using OASIS 2 [68] with *p* values calculated by the log-rank (Mantel–Cox) test and multiple corrections applied using Bonferroni procedure. Graphs were plotted using GraphPad Prism v9.

### Microscopy and fluorescence quantitation

GFP expression was imaged using a Leica DM5500B compound scope with Leica (LAS X) imaging software and quantified using observational scoring of populations. Progeny of transgenic mothers in different genetic backgrounds were grown to the late larval L4 stage then examined for fluorescence levels after 12 h (early day 1 adults) and 24 h (late day 1 adults). Animals were immobilized using Levamisole (10 mM) and imaged under set magnification and intensity settings, until ≥15 animals were imaged per condition.

### RNA isolation

Animals were synchronized by bleach treatment and hatched in M9 until arrested as L1 larvae. Synchronized larvae were plated on NGM plates containing OP50 and grown to gravid adult stage (72 hours post L1 synchronization). Animals were washed three times in M9 buffer, flash frozen in liquid nitrogen, and lysed in Trizol. RNA was isolated using two rounds of chloroform extraction followed by isopropanol precipitation.

### sRNA-seq library preparation

16-30-nt RNAs were size selected from total RNA on 17% denaturing polyacrylamide gels. Small RNAs were treated with RNA polyphosphatase to reduce 5′ di- and triphosphates to monophosphates to enable 5′ adapter ligation. Sequencing libraries were prepared with the NEBNext Multiplex Small RNA Library Prep Set for Illumina (NEB, Cat # E7300S). Libraries were size selected on 10% polyacrylamide gels and sequenced on an Illumina NextSeq 500 sequencer (High Output Kit, Single-End, 75 Cycles).

### sRNA-seq data analysis

sRNA sequencing data was processed using the tinyRNA pipeline with default settings and *C. elegans* WS279 genome release [71–75]. 22G-RNA and other small RNA annotations utilized GFF3-formatted annotations available from Knittel et al [76]. 22G-RNAs were defined as antisense-strand 21-23-nt reads containing the 5′-G hallmark of this class of small RNA. Differential expression analysis was done with DESeq2 within the tinyRNA pipeline using the Wald test significance assessment [77]. *p* values were adjusted for multiple testing using the Benjamini-Hochberg procedure. Plotting was done using Matplotlib, R, IGV, and Adobe Illustrator [78–80].

### mRNA-seq library preparation

rRNA was depleted using the Ribo-Zero rRNA Removal Kit (Human/Mouse/Rat, Illumina Cat # MRZH11124). rRNA-depleted RNA was DNase treated and size selected (>200 nt) to remove 5S rRNA and tRNA using RNA the Clean & Concentrator-5 Kit (Zymo Research, Cat # R1015). RNA-seq libraries were prepared using the NEBNext Ultra II Directional RNA Library Prep Kit for Illumina (NEB, Cat # E7760S). Samples were sequenced on an Illumina HiSeq (Paired-End, 150 Cycles).

### mRNA-seq data analysis

fastp was used to remove adapters and filter low quality data (fastp -w 16 -q 30 -u 70 -l 30 -r -W 4 -M 20) [72]. Reads were mapped to the *C. elegans* genome (Wormbase release WS279) and transcripts were quantified using RSEM v1.3.1 with STAR v2.7.10b as the aligner [81, 82]. Differential expression analysis was done using DESeq2 v1.50.2 [77].

### Integrative analysis of sRNA and mRNA data

Integrative analysis of sRNA- and mRNA-seq data was done using the RNA-integrate pipeline (available from https://github.com/MontgomeryLab/RNA-integrate). Within the pipeline, differential expression analysis of sRNA-seq and mRNA-seq data were done in parallel with DESeq2 v1.50.2 using the counts tables from the tinyRNA and RSEM analyses described above [75, 77, 82]. Changes in WAGO 22G-RNAs levels were compared to changes in the corresponding mRNAs in *tcer-1* mutants relative to wildtype and data was plotted in Plotly.js v2.11.1 based on fold change, read abundance, and *p* value, as indicated in the plots.

## Acknowledgements

The authors are grateful to Dr. Mayur Devare for assistance with data presentation. Thanks to members of the Ghazi and Montgomery labs and the Pittsburgh ‘Wormclub’ community for valuable inputs throughout this study. Some strains were provided by the CGC, which is funded by NIH Office of Research Infrastructure Programs (P40 OD010440).

## Data Availability

Raw and processed high-throughput sequencing data generated in this study are available from the NCBI Gene Expression Omnibus (GEO) under accession number GSE316795.

## List of Supplementary Material

**Supplementay Figure 1**: Survival of *tcer-1(glm27)* mutants on PA14.

**Table S1.** Impact of *tcer-1* RNAi on survival of *ppw-1* and *rrf-1* mutants on PA14.

**Table S2.** Impact of *ppw-1* and *rrf-1* mutations on survival of *tcer-1* mutants on PA14.

**Table S3.** Impact of loss of small RNA biogenesis factors on worm survival on PA14.

**Table S4.** Differential expression analysis of small RNAs in *tcer-1* mutants vs wildtype.

**Table S5.** Geometric mean-normalized counts and differential expression analysis of mRNA in *tcer-1* mutants vs wild type.

**Table S6.** Integrative analysis of 22G-RNA and WAGO target mRNA expression in *tcer-1(tm1452)* vs wildtype.

**Table S7.** Integrative analysis of 22G-RNA and WAGO target mRNA expression in *tcer-1(glm27)* vs wildtype.

## REFERENCES

1. Jouravleva K, Zamore PD. A guide to the biogenesis and functions of endogenous small non-coding RNAs in animals. Nat Rev Mol Cell Biol. 2025;26(5):347–70. Epub 20250124. doi: 10.1038/s41580-024-00818-9. PubMed PMID: 39856370.

2. Ketting RF. The many faces of RNAi. Dev Cell. 2011;20(2):148–61. doi: 10.1016/j.devcel.2011.01.012. PubMed PMID: 21316584.

3. Bernier A, Sagan SM. The Diverse Roles of microRNAs at the Host⁻Virus Interface. Viruses. 2018;10(8). Epub 20180819. doi: 10.3390/v10080440. PubMed PMID: 30126238; PubMed Central PMCID: PMCPMC6116274.

4. Mehta A, Baltimore D. MicroRNAs as regulatory elements in immune system logic. Nat Rev Immunol. 2016;16(5):279–94. doi: 10.1038/nri.2016.40. PubMed PMID: 27121651.

5. Nejad C, Stunden HJ, Gantier MP. A guide to miRNAs in inflammation and innate immune responses. Febs j. 2018;285(20):3695–716. Epub 20180506. doi: 10.1111/febs.14482. PubMed PMID: 29688631.

6. Xiao C, Rajewsky K. MicroRNA control in the immune system: basic principles. Cell. 2009;136(1):26–36. doi: 10.1016/j.cell.2008.12.027. PubMed PMID: 19135886.

7. Yan T, Lu R. Shared and unique mechanisms of RNAi-mediated antiviral immunity in C. elegans. Virology. 2025;605:110459. Epub 20250221. doi: 10.1016/j.virol.2025.110459. PubMed PMID: 40022946; PubMed Central PMCID: PMCPMC11970214.

8. Gammon DB, Mello CC. RNA interference-mediated antiviral defense in insects. Current Opinion in Insect Science. 2015;8:111–20. doi: 10.1016/j.cois.2015.01.006.

9. Huang CY, Wang H, Hu P, Hamby R, Jin H. Small RNAs - Big Players in Plant-Microbe Interactions. Cell Host Microbe. 2019;26(2):173–82. doi: 10.1016/j.chom.2019.07.021. PubMed PMID: 31415750.

10. Yang Z, Li Y. Dissection of RNAi-based antiviral immunity in plants. Curr Opin Virol. 2018;32:88–99. Epub 20181031. doi: 10.1016/j.coviro.2018.08.003. PubMed PMID: 30388659.

11. Berkhout B. RNAi-mediated antiviral immunity in mammals. Curr Opin Virol. 2018;32:9–14. Epub 20180714. doi: 10.1016/j.coviro.2018.07.008. PubMed PMID: 30015014.

12. Cullen BR, Cherry S, tenOever BR. Is RNA interference a physiologically relevant innate antiviral immune response in mammals? Cell Host Microbe. 2013;14(4):374–8. doi: 10.1016/j.chom.2013.09.011. PubMed PMID: 24139396.

13. Svoboda P. Renaissance of mammalian endogenous RNAi. FEBS Letters. 2014;588(15):2550–6. doi: 10.1016/j.febslet.2014.05.030.

14. Takahashi T, Heaton SM, Parrish NF. Mammalian antiviral systems directed by small RNA. PLoS Pathog. 2021;17(12):e1010091. Epub 20211216. doi: 10.1371/journal.ppat.1010091. PubMed PMID: 34914813; PubMed Central PMCID: PMCPMC8675686.

15. Abbas MN, Kausar S, Asma B, Ran W, Li J, Lin Z, et al. MicroRNAs reshape the immunity of insects in response to bacterial infection. Front Immunol. 2023;14:1176966. Epub 20230421. doi: 10.3389/fimmu.2023.1176966. PubMed PMID: 37153604; PubMed Central PMCID: PMCPMC10161253.

16. Eulalio A, Schulte L, Vogel J. The mammalian microRNA response to bacterial infections. RNA Biol. 2012;9(6):742–50. Epub 20120601. doi: 10.4161/rna.20018. PubMed PMID: 22664920.

17. Jin H. Endogenous small RNAs and antibacterial immunity in plants. FEBS Lett. 2008;582(18):2679–84. Epub 20080711. doi: 10.1016/j.febslet.2008.06.053. PubMed PMID: 18619960; PubMed Central PMCID: PMCPMC5912937.

18. Kudlow BA, Zhang L, Han M. Systematic analysis of tissue-restricted miRISCs reveals a broad role for microRNAs in suppressing basal activity of the C. elegans pathogen response. Mol Cell. 2012;46(4):530–41. Epub 20120411. doi: 10.1016/j.molcel.2012.03.011. PubMed PMID: 22503424; PubMed Central PMCID: PMCPMC3365535.

19. Martineau CN, Kirienko NV, Pujol N. Innate immunity in C. elegans. Curr Top Dev Biol. 2021;144:309–51. Epub 20210304. doi: 10.1016/bs.ctdb.2020.12.007. PubMed PMID: 33992157; PubMed Central PMCID: PMCPMC9175240.

20. Charlesworth AG, Seroussi U, Lehrbach NJ, Renaud MS, Sundby AE, Molnar RI, et al. Two isoforms of the essential C. elegans Argonaute CSR-1 differentially regulate sperm and oocyte fertility. Nucleic Acids Res. 2021;49(15):8836–65. doi: 10.1093/nar/gkab619. PubMed PMID: 34329465; PubMed Central PMCID: PMCPMC8421154.

21. Dai LL, Gao JX, Zou CG, Ma YC, Zhang KQ. mir-233 modulates the unfolded protein response in C. elegans during Pseudomonas aeruginosa infection. PLoS Pathog. 2015;11(1):e1004606. Epub 20150108. doi: 10.1371/journal.ppat.1004606. PubMed PMID: 25569229; PubMed Central PMCID: PMCPMC4287614.

22. Ren Z, Ambros VR. Caenorhabditis elegans microRNAs of the let-7 family act in innate immune response circuits and confer robust developmental timing against pathogen stress. Proc Natl Acad Sci U S A. 2015;112(18):E2366–75. Epub 20150420. doi: 10.1073/pnas.1422858112. PubMed PMID: 25897023; PubMed Central PMCID: PMCPMC4426397.

23. Seroussi U, Lugowski A, Wadi L, Lao RX, Willis AR, Zhao W, et al. A comprehensive survey of C. elegans argonaute proteins reveals organism-wide gene regulatory networks and functions. Elife. 2023;12. Epub 20230215. doi: 10.7554/eLife.83853. PubMed PMID: 36790166; PubMed Central PMCID: PMCPMC10101689.

24. Sun L, Zhi L, Shakoor S, Liao K, Wang D. microRNAs Involved in the Control of Innate Immunity in Candida Infected Caenorhabditis elegans. Sci Rep. 2016;6:36036. Epub 20161031. doi: 10.1038/srep36036. PubMed PMID: 27796366; PubMed Central PMCID: PMCPMC5086856.

25. Zhi L, Yu Y, Li X, Wang D, Wang D. Molecular Control of Innate Immune Response to Pseudomonas aeruginosa Infection by Intestinal let-7 in Caenorhabditis elegans. PLoS Pathog. 2017;13(1):e1006152. Epub 20170117. doi: 10.1371/journal.ppat.1006152. PubMed PMID: 28095464; PubMed Central PMCID: PMCPMC5271417.

26. Ketting RF, Cochella L. Concepts and functions of small RNA pathways in C. elegans. Curr Top Dev Biol. 2021;144:45–89. Epub 20201003. doi: 10.1016/bs.ctdb.2020.08.002. PubMed PMID: 33992161.

27. Kaletsky R, Moore RS, Vrla GD, Parsons LR, Gitai Z, Murphy CT. C. elegans interprets bacterial non-coding RNAs to learn pathogenic avoidance. Nature. 2020;586(7829):445–51. Epub 20200909. doi: 10.1038/s41586-020-2699-5. PubMed PMID: 32908307; PubMed Central PMCID: PMCPMC8547118.

28. Moore RS, Kaletsky R, Lesnik C, Cota V, Blackman E, Parsons LR, et al. The role of the Cer1 transposon in horizontal transfer of transgenerational memory. Cell. 2021;184(18):4697–712 e18. Epub 20210806. doi: 10.1016/j.cell.2021.07.022. PubMed PMID: 34363756; PubMed Central PMCID: PMCPMC8812995.

29. Moore RS, Kaletsky R, Murphy CT. Piwi/PRG-1 Argonaute and TGF-beta Mediate Transgenerational Learned Pathogenic Avoidance. Cell. 2019;177(7):1827–41 e12. Epub 20190606. doi: 10.1016/j.cell.2019.05.024. PubMed PMID: 31178117; PubMed Central PMCID: PMCPMC7518193.

30. Montes M, Coiras M, Becerra S, Moreno-Castro C, Mateos E, Majuelos J, et al. Functional Consequences for Apoptosis by Transcription Elongation Regulator 1 (TCERG1)-Mediated Bcl-x and Fas/CD95 Alternative Splicing. PLoS One. 2015;10(10):e0139812. doi: 10.1371/journal.pone.0139812. PubMed PMID: 26462236; PubMed Central PMCID: PMCPMC4604205.

31. Moreno-Castro C, Prieto-Sánchez S, Sánchez-Hernández N, Hernández-Munain C, Suñé C. Role for the splicing factor TCERG1 in Cajal body integrity and snRNP assembly. Journal of Cell Science. 2019;132(22). doi: 10.1242/jcs.232728.

32. Munoz-Cobo JP, Sanchez-Hernandez N, Gutierrez S, El Yousfi Y, Montes M, Gallego C, et al. Transcriptional Elongation Regulator 1 Affects Transcription and Splicing of Genes Associated with Cellular Morphology and Cytoskeleton Dynamics and Is Required for Neurite Outgrowth in Neuroblastoma Cells and Primary Neuronal Cultures. Mol Neurobiol. 2017;54(10):7808–23. doi: 10.1007/s12035-016-0284-6. PubMed PMID: 27844289.

33. Pearson JL, Robinson TJ, Munoz MJ, Kornblihtt AR, Garcia-Blanco MA. Identification of the cellular targets of the transcription factor TCERG1 reveals a prevalent role in mRNA processing. J Biol Chem. 2008;283(12):7949–61. doi: 10.1074/jbc.M709402200. PubMed PMID: 18187414.

34. Sanchez-Hernandez N, Boireau S, Schmidt U, Munoz-Cobo JP, Hernandez-Munain C, Bertrand E, et al. The in vivo dynamics of TCERG1, a factor that couples transcriptional elongation with splicing. RNA. 2016;22(4):571–82. doi: 10.1261/rna.052795.115. PubMed PMID: 26873599; PubMed Central PMCID: PMCPMC4793212.

35. Amrit FRG, Naim N, Ratnappan R, Loose J, Mason C, Steenberge L, et al. The Longevity-Promoting Factor, TCER-1, Widely Represses Stress Resistance and Innate Immunity. Nature Communications. 2019;10(1):3042. Epub 2019/07/19. doi: 10.1038/s41467-019-10759-z. PubMed PMID: 31316054; PubMed Central PMCID: PMCPMC6637209.

36. Hernando CE, García Hourquet M, de Leone MJ, Careno D, Iserte J, Mora Garcia S, et al. A Role for Pre-mRNA-PROCESSING PROTEIN 40C in the Control of Growth, Development, and Stress Tolerance in Arabidopsis thaliana. Frontiers in plant science. 2019;10:1019–. doi: 10.3389/fpls.2019.01019. PubMed PMID: 31456814.

37. Kim JK, Gabel HW, Kamath RS, Tewari M, Pasquinelli A, Rual JF, et al. Functional genomic analysis of RNA interference in C. elegans. Science. 2005;308(5725):1164–7. Epub 20050324. doi: 10.1126/science.1109267. PubMed PMID: 15790806.

38. Montgomery TA, Rim YS, Zhang C, Dowen RH, Phillips CM, Fischer SE, et al. PIWI associated siRNAs and piRNAs specifically require the Caenorhabditis elegans HEN1 ortholog henn-1. PLoS Genet. 2012;8(4):e1002616. Epub 20120419. doi: 10.1371/journal.pgen.1002616. PubMed PMID: 22536158; PubMed Central PMCID: PMCPMC3334881.

39. Pushpa K, Kumar GA, Subramaniam K. PUF-8 and TCER-1 are essential for normal levels of multiple mRNAs in the C. elegans germline. Development. 2013;140(6):1312–20. doi: 10.1242/dev.087833. PubMed PMID: 23444359; PubMed Central PMCID: PMCPMC3585663.

40. Kumsta C, Hansen M. C. elegans rrf-1 mutations maintain RNAi efficiency in the soma in addition to the germline. PLoS One. 2012;7(5):e35428. Epub 20120504. doi: 10.1371/journal.pone.0035428. PubMed PMID: 22574120; PubMed Central PMCID: PMCPMC3344830.

41. Tijsterman M, Okihara KL, Thijssen K, Plasterk RH. PPW-1, a PAZ/PIWI protein required for efficient germline RNAi, is defective in a natural isolate of C. elegans. Curr Biol. 2002;12(17):1535–40. doi: 10.1016/s0960-9822(02)01110-7. PubMed PMID: 12225671.

42. Ketting RF, Haverkamp TH, van Luenen HG, Plasterk RH. Mut-7 of C. elegans, required for transposon silencing and RNA interference, is a homolog of Werner syndrome helicase and RNaseD. Cell. 1999;99(2):133–41. doi: 10.1016/s0092-8674(00)81645-1. PubMed PMID: 10535732.

43. Phillips Carolyn M, Montgomery Brooke E, Breen Peter C, Roovers Elke F, Rim Y-S, Ohsumi Toshiro K, et al. MUT-14 and SMUT-1 DEAD Box RNA Helicases Have Overlapping Roles in Germline RNAi and Endogenous siRNA Formation. Current Biology. 2014;24(8):839–44. doi: 10.1016/j.cub.2014.02.060.

44. Phillips CM, Montgomery TA, Breen PC, Ruvkun G. MUT-16 promotes formation of perinuclear mutator foci required for RNA silencing in the C. elegans germline. Genes Dev. 2012;26(13):1433–44. Epub 2012/06/21. doi: 10.1101/gad.193904.112. PubMed PMID: 22713602; PubMed Central PMCID: PMCPMC3403012.

45. Tijsterman M, Ketting RF, Okihara KL, Sijen T, Plasterk RH. RNA helicase MUT-14-dependent gene silencing triggered in C. elegans by short antisense RNAs. Science. 2002;295(5555):694–7. doi: 10.1126/science.1067534. PubMed PMID: 11809977.

46. Vastenhouw NL, Fischer SE, Robert VJ, Thijssen KL, Fraser AG, Kamath RS, et al. A genome-wide screen identifies 27 genes involved in transposon silencing in C. elegans. Curr Biol. 2003;13(15):1311–6. doi: 10.1016/s0960-9822(03)00539-6. PubMed PMID: 12906791.

47. Chen CC, Simard MJ, Tabara H, Brownell DR, McCollough JA, Mello CC. A member of the polymerase beta nucleotidyltransferase superfamily is required for RNA interference in C. elegans. Curr Biol. 2005;15(4):378–83. doi: 10.1016/j.cub.2005.01.009. PubMed PMID: 15723801.

48. Shukla A, Yan J, Pagano DJ, Dodson AE, Fei Y, Gorham J, et al. poly(UG)-tailed RNAs in genome protection and epigenetic inheritance. Nature. 2020;582(7811):283–8. Epub 20200520. doi: 10.1038/s41586-020-2323-8. PubMed PMID: 32499657; PubMed Central PMCID: PMCPMC8396162.

49. Schott DH, Cureton DK, Whelan SP, Hunter CP. An antiviral role for the RNA interference machinery in Caenorhabditis elegans. Proc Natl Acad Sci U S A. 2005;102(51):18420–4. Epub 20051209. doi: 10.1073/pnas.0507123102. PubMed PMID: 16339901; PubMed Central PMCID: PMCPMC1317933.

50. Tabara H, Sarkissian M, Kelly WG, Fleenor J, Grishok A, Timmons L, et al. The rde-1 gene, RNA interference, and transposon silencing in C. elegans. Cell. 1999;99(2):123–32. doi: 10.1016/s0092-8674(00)81644-x. PubMed PMID: 10535731.

51. Gu W, Shirayama M, Conte D, Jr., Vasale J, Batista PJ, Claycomb JM, et al. Distinct argonaute-mediated 22G-RNA pathways direct genome surveillance in the C. elegans germline. Mol Cell. 2009;36(2):231–44. Epub 20091001. doi: 10.1016/j.molcel.2009.09.020. PubMed PMID: 19800275; PubMed Central PMCID: PMCPMC2776052.

52. Ashe A, Belicard T, Le Pen J, Sarkies P, Frezal L, Lehrbach NJ, et al. A deletion polymorphism in the Caenorhabditis elegans RIG-I homolog disables viral RNA dicing and antiviral immunity. Elife. 2013;2:e00994. Epub 20131008. doi: 10.7554/eLife.00994. PubMed PMID: 24137537; PubMed Central PMCID: PMCPMC3793227.

53. Lu R, Maduro M, Li F, Li HW, Broitman-Maduro G, Li WX, et al. Animal virus replication and RNAi-mediated antiviral silencing in Caenorhabditis elegans. Nature. 2005;436(7053):1040–3. doi: 10.1038/nature03870. PubMed PMID: 16107851; PubMed Central PMCID: PMCPMC1388260.

54. Wilkins C, Dishongh R, Moore SC, Whitt MA, Chow M, Machaca K. RNA interference is an antiviral defence mechanism in Caenorhabditis elegans. Nature. 2005;436(7053):1044–7. doi: 10.1038/nature03957. PubMed PMID: 16107852.

55. Felix MA, Ashe A, Piffaretti J, Wu G, Nuez I, Belicard T, et al. Natural and experimental infection of Caenorhabditis nematodes by novel viruses related to nodaviruses. PLoS Biol. 2011;9(1):e1000586. Epub 20110125. doi: 10.1371/journal.pbio.1000586. PubMed PMID: 21283608; PubMed Central PMCID: PMCPMC3026760.

56. Knittel TL, Montgomery BE, Sprister RA, Magelky CN, Smith MJ, Soto-Ojeda M, et al. Argonaute-siRNA loading via the RNA-binding protein RDE-4 in C. elegans. Curr Biol. 2025;35(23):5897–907 e6. Epub 20251118. doi: 10.1016/j.cub.2025.10.042. PubMed PMID: 41260219; PubMed Central PMCID: PMCPMC12638004.

57. Levin M, Hashimshony T, Wagner F, Yanai I. Developmental milestones punctuate gene expression in the Caenorhabditis embryo. Dev Cell. 2012;22(5):1101–8. Epub 20120503. doi: 10.1016/j.devcel.2012.04.004. PubMed PMID: 22560298.

58. Reed KJ, Svendsen JM, Brown KC, Montgomery BE, Marks TN, Vijayasarathy T, et al. Widespread roles for piRNAs and WAGO-class siRNAs in shaping the germline transcriptome of Caenorhabditis elegans. Nucleic Acids Res. 2020;48(4):1811–27. doi: 10.1093/nar/gkz1178. PubMed PMID: 31872227; PubMed Central PMCID: PMCPMC7038979.

59. Seistrup A-S, Nischwitz E, Butter F, Ketting RF. Crosstalk between and developmental dynamics of C. elegans Argonaute proteins. bioRxiv. 2025:2025.12.17.694999. doi: 10.64898/2025.12.17.694999.

60. He Y, Liu J, Grossman D, Durrant D, Sweatman T, Lothstein L, et al. Phosphorylation of mitochondrial phospholipid scramblase 3 by protein kinase C-delta induces its activation and facilitates mitochondrial targeting of tBid. J Cell Biochem. 2007;101(5):1210–21. doi: 10.1002/jcb.21243. PubMed PMID: 17226776.

61. Liu J, Dai Q, Chen J, Durrant D, Freeman A, Liu T, et al. Phospholipid scramblase 3 controls mitochondrial structure, function, and apoptotic response. Mol Cancer Res. 2003;1(12):892–902. PubMed PMID: 14573790.

62. Liu J, Epand RF, Durrant D, Grossman D, Chi NW, Epand RM, et al. Role of phospholipid scramblase 3 in the regulation of tumor necrosis factor-alpha-induced apoptosis. Biochemistry. 2008;47(15):4518–29. Epub 20080322. doi: 10.1021/bi701962c. PubMed PMID: 18358005.

63. Anderson SM, Pukkila-Worley R. Immunometabolism in Caenorhabditis elegans. PLoS Pathog. 2020;16(10):e1008897. Epub 20201008. doi: 10.1371/journal.ppat.1008897. PubMed PMID: 33031414; PubMed Central PMCID: PMCPMC7544110.

64. Bahr L, Amrit FRG, Silvia PE, Wayhs B, Osman GA, Devare MN, et al. LIPL-1 and LIPL-2 are TCER-1-regulated lysosomal lipases with distinct roles in immunity and fertility. PLoS Genet. 2025;21(12):e1011804. Epub 20251212. doi: 10.1371/journal.pgen.1011804. PubMed PMID: 41385585; PubMed Central PMCID: PMCPMC12716718.

65. Yigit E, Batista PJ, Bei Y, Pang KM, Chen CC, Tolia NH, et al. Analysis of the C. elegans Argonaute family reveals that distinct Argonautes act sequentially during RNAi. Cell. 2006;127(4):747–57. doi: 10.1016/j.cell.2006.09.033. PubMed PMID: 17110334.

66. Wang C, Sheng W, Zhou Y, Hang X, Zhao J, Gu Y, et al. siRNA-AGO2 complex inhibits bacterial gene translation: A promising therapeutic strategy for superbug infection. Cell Rep Med. 2025;6(3):101997. Epub 20250306. doi: 10.1016/j.xcrm.2025.101997. PubMed PMID: 40054457; PubMed Central PMCID: PMCPMC11970400.

67. Brenner S. The genetics of Caenorhabditis elegans. Genetics. 1974;77(1):71–94. Epub 1974/05/01. doi: 10.1093/genetics/77.1.71. PubMed PMID: 4366476; PubMed Central PMCID: PMCPMC1213120.

68. Han SK, Lee D, Lee H, Kim D, Son HG, Yang JS, et al. OASIS 2: online application for survival analysis 2 with features for the analysis of maximal lifespan and healthspan in aging research. Oncotarget. 2016;7(35):56147–52. doi: 10.18632/oncotarget.11269. PubMed PMID: 27528229; PubMed Central PMCID: PMCPMC5302902.

69. Keith SA, Amrit FR, Ratnappan R, Ghazi A. The C. elegans healthspan and stress-resistance assay toolkit. Methods. 2014;68(3):476–86. Epub 20140413. doi: 10.1016/j.ymeth.2014.04.003. PubMed PMID: 24727065.

70. Tan MW, Ausubel FM. Caenorhabditis elegans: a model genetic host to study Pseudomonas aeruginosa pathogenesis. Curr Opin Microbiol. 2000;3(1):29–34. doi: 10.1016/s1369-5274(99)00047-8. PubMed PMID: 10679415.

71. Anders S, Pyl PT, Huber W. HTSeq--a Python framework to work with high-throughput sequencing data. Bioinformatics. 2015;31(2):166–9. Epub 20140925. doi: 10.1093/bioinformatics/btu638. PubMed PMID: 25260700; PubMed Central PMCID: PMCPMC4287950.

72. Chen S, Zhou Y, Chen Y, Gu J. fastp: an ultra-fast all-in-one FASTQ preprocessor. Bioinformatics. 2018;34(17):i884–i90. doi: 10.1093/bioinformatics/bty560. PubMed PMID: 30423086; PubMed Central PMCID: PMCPMC6129281.

73. Langmead B, Trapnell C, Pop M, Salzberg SL. Ultrafast and memory-efficient alignment of short DNA sequences to the human genome. Genome Biol. 2009;10(3):R25. Epub 20090304. doi: 10.1186/gb-2009-10-3-r25. PubMed PMID: 19261174; PubMed Central PMCID: PMCPMC2690996.

74. Sternberg PW, Van Auken K, Wang Q, Wright A, Yook K, Zarowiecki M, et al. WormBase 2024: status and transitioning to Alliance infrastructure. Genetics. 2024;227(1). doi: 10.1093/genetics/iyae050. PubMed PMID: 38573366; PubMed Central PMCID: PMCPMC11075546.

75. Tate AJ, Brown KC, Montgomery TA. tiny-count: a counting tool for hierarchical classification and quantification of small RNA-seq reads with single-nucleotide precision. Bioinform Adv. 2023;3(1):vbad065. Epub 20230518. doi: 10.1093/bioadv/vbad065. PubMed PMID: 37288323; PubMed Central PMCID: PMCPMC10243934.

76. Knittel TL, Montgomery BE, Tate AJ, Deihl EW, Nawrocki AS, Hoerndli FJ, et al. A low-abundance class of Dicer-dependent siRNAs produced from a variety of features in C. elegans. Genome Res. 2024;34(12):2203–16. Epub 20241223. doi: 10.1101/gr.279083.124. PubMed PMID: 39622635; PubMed Central PMCID: PMCPMC11694761.

77. Love MI, Huber W, Anders S. Moderated estimation of fold change and dispersion for RNA-seq data with DESeq2. Genome Biol. 2014;15(12):550. doi: 10.1186/s13059-014-0550-8. PubMed PMID: 25516281; PubMed Central PMCID: PMCPMC4302049.

78. Hunter JD. Matplotlib: A 2D Graphics Environment. Computing in Science & Engineering. 2007;9(3):90–5. doi: 10.1109/MCSE.2007.55.

79. Robinson JT, Thorvaldsdóttir H, Winckler W, Guttman M, Lander ES, Getz G, et al. Integrative genomics viewer. Nat Biotechnol. 2011;29(1):24–6. doi: 10.1038/nbt.1754. PubMed PMID: 21221095; PubMed Central PMCID: PMCPMC3346182.

80. R Core Team. R: a Language and Environment for Statistical Computing. R Foundation for Statistical Computing, Vienna, Austria. 2021.

81. Dobin A, Davis CA, Schlesinger F, Drenkow J, Zaleski C, Jha S, et al. STAR: ultrafast universal RNA-seq aligner. Bioinformatics. 2013;29(1):15–21. Epub 20121025. doi: 10.1093/bioinformatics/bts635. PubMed PMID: 23104886; PubMed Central PMCID: PMCPMC3530905.

82. Li B, Dewey CN. RSEM: accurate transcript quantification from RNA-Seq data with or without a reference genome. BMC Bioinformatics. 2011;12:323. Epub 20110804. doi: 10.1186/1471-2105-12-323. PubMed PMID: 21816040; PubMed Central PMCID: PMCPMC3163565.

