## Supplementary Material- Fig S1, Tables S1-S3 for "A TCER-1-siRNA Regulatory Axis Suppresses Antibacterial Innate Immunity in *C. elegans*"

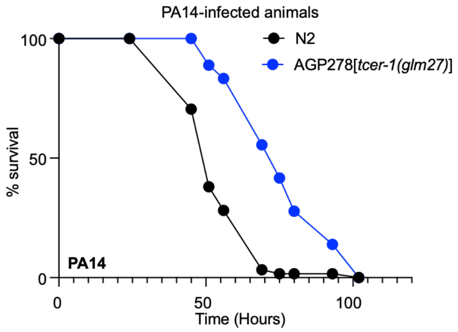


| Survival of *tcer-1(glm27)* mutant upon PA14 exposure | | | | | |
|---|---|---|---|---|---|
| **Strain** | **Genotype** | **n = obs/ total** | **Mean (hrs)** | **SEM** | **p (vs. N2 ctrl)** |
| **Trial #1** | | | | | |
| N2 | Wildtype | 90 | 55.43 | 1.32 | |
| AGP278a - Line #1 | *tcer-1(glm27)* | 48 | 90.03 | 4.26 | 0.001 |
| AGP278a - Line #2 | *tcer-1(glm27)* | 30 | 83.05 | 5.5 | 8.70E-08 |
| **Trial #2** | | | | | |
| N2 | Wildtype | 120 | 52.56 | 0.94 | |
| AGP278a - Line #1 | *tcer-1(glm27)* | 23 | 73.45 | 5.08 | 0.00000015 |
| AGP278a - Line #2 | *tcer-1(glm27)* | 61 | 83.76 | 4.29 | 0.00E+00 |

**Supplementary Figure S1:** Survival of *tcer-1(glm27)* (blue) mutants following PA14 infection compared to WT animals (black). Data from two independent trials is summarized in the table showing mean survival in hours (mean) and standard error from the mean (SEM). n=observed/total (see Methods for details). Data from Trial 2 is plotted in the graph. *p* values were calculated using the log-rank method (Mantel Cox).

**Table S1.** Impact of *tcer-1* RNAi on survival of *ppw-1* and *rrf-1* mutants on PA14

| Genotype | RNAi | n = obs/total | Mean (hrs) | SEM | *p* (vs. Mutant pAD12 Ctrl) | *p* (vs. N2 pAD12 Ctrl) |
|---|---|---|---|---|---|---|
| **Trial 1** | | | | | | |
| N2 | Ctrl | 87/113 | 56.61 | 1.36 | | |
| N2 | *tcer-1* | 86/114 | 64.57 | 1.51 | | 0.0007 |
| *rrf-1(ok589)* | Ctrl | 97/117 | 75.99 | 2.25 | | <0.0001 |
| *rrf-1(ok589)* | *tcer-1* | 119/136 | 83.89 | 1.75 | 0.3887 | |
| *ppw-1(pk1425)* | Ctrl | 118/126 | 76.33 | 1.77 | | <0.0001 |
| *ppw-1(pk1425)* | *tcer-1* | 132/148 | 70 | 1.32 | 0.0197 | |
| **Trial 2** | | | | | | |
| N2 | Ctrl | 110/134 | 60.93 | 0.95 | | |
| N2 | *tcer-1* | 129/157 | 67.91 | 1.09 | 0.00004 | |
| *rrf-1(ok589)* | Ctrl | 114/136 | 77.45 | 1.67 | <0.0001 | |
| *rrf-1(ok589)* | *tcer-1* | 117/154 | 77.3 | 1.33 | | 1 |
| *ppw-1(pk1425)* | Ctrl | 123/144 | 72.14 | 1.46 | <0.0001 | |
| *ppw-1(pk1425)* | *tcer-1* | 1412/154 | 72.46 | 1.2 | | 1 |
| **Trial 3** | | | | | | |
| N2 | Ctrl | 63/81 | 58.52 | 1.48 | | |
| N2 | *tcer-1* | 78/109 | 69.23 | 1.48 | | 0.0000068 |
| *rrf-1(ok589)* | Ctrl | 93/108 | 83.39 | 2.16 | | <0.0001 |
| *rrf-1(ok589)* | *tcer-1* | 104/131 | 88.44 | 2.12 | 0.3968 | |
| *ppw-1(pk1425)* | Ctrl | 99/112 | 71.2 | 1.95 | | 0.000007 |
| *ppw-1(pk1425)* | *tcer-1* | 110/121 | 70.14 | 1.62 | 1 | |
| **Trial 4** | | | | | | |
| N2 | Ctrl | 131/167 | 59.33 | 1.16 | | |
| N2 | *tcer-1* | 143/161 | 76.23 | 1.52 | | <0.0001 |
| *rrf-1(ok589)* | Ctrl | 137/170 | 79.77 | 1.63 | | <0.0001 |
| *rrf-1(ok589)* | *tcer-1* | 144/164 | 82.62 | 1.69 | 0.807 | |
| *ppw-1(pk1425)* | Ctrl | 144/157 | 84.43 | 1.74 | | <0.0001 |
| *ppw-1(pk1425)* | *tcer-1* | 134/151 | 87.16 | 1.99 | 1 | |
| **Trial 5** | | | | | | |
| N2 | Ctrl | 75/151 | 57.93 | 1.09 | | |
| N2 | *tcer-1* | 73/175 | 77.06 | 1.8 | | 0.0001 |
| *rrf-1(ok589)* | Ctrl | 97/155 | 81.81 | 2.1 | | 0.0001 |
| *rrf-1(ok589)* | *tcer-1* | 123/160 | 82.11 | 1.24 | 0.2123 | |
| *ppw-1(pk1425)* | Ctrl | 71/140 | 70.34 | 2.11 | | 0.000002 |
| *ppw-1(pk1425)* | *tcer-1* | 111/155 | 78.32 | 1.32 | 0.0019 | |
| **Trial 6** | | | | | | |
| N2 | Ctrl | 99/112 | 64.21 | 1.73 | | |
| N2 | *tcer-1* | 92/97 | 78.73 | 1.39 | | <0.0001 |
| *rrf-1(ok589)* | Ctrl | 100/115 | 78.17 | 2.8 | | 0.0001 |
| *rrf-1(ok589)* | *tcer-1* | 126/140 | 93.07 | 2.4 | 0.0076 | |
| *ppw-1(pk1425)* | Ctrl | 82/123 | 86.34 | 2.54 | | <0.0001 |
| *ppw-1(pk1425)* | *tcer-1* | 149/180 | 91.81 | 1.83 | 0.6764 | |

**Table S2.** Impact of loss of *tcer-1* on survival of *ppw-1* and *rrf-1* mutants on PA14

| Strain | Genotype | n = obs/total | Mean (hrs) | SEM | p (vs N2) | *p* (vs *tcer-1*) |
|---|---|---|---|---|---|---|
| **Trial 1** | | | | | | |
| N2 | Wildtype | 108/130 | 74.9 | 1.34 | | |
| CF1266 | *tcer-1(tm1452)* | 91/120 | 103.91 | 2.45 | <0.0001 | |
| NL3511 | *ppw-1(pk1425)* | 119/146 | 97.77 | 1.96 | <0.0001 | |
| AGP274 | *ppw-1; tcer-1* | 99/123 | 112.97 | 2.94 | <0.0001 | 0.05 |
| RB798 | *rrf-1(ok589)* | 113/137 | 97.99 | 1.89 | <0.0001 | |
| AGP273 | *rrf-1;tcer-1* | 88/106 | 108.5 | 2.55 | <0.0001 | 0.63 |
| **Trial 2** | | | | | | |
| N2 | Wildtype | 116/143 | 88.97 | 2.01 | | |
| CF1266 | *tcer-1(tm1452)* | 96/111 | 102.76 | 2.3 | 0.0002 | |
| NL3511 | *ppw-1(pk1425)* | 134/148 | 102.95 | 2.38 | 0.0001 | |
| AGP274 | *ppw-1; tcer-1* | 95/116 | 106.68 | 2.94 | <0.0001 | 0.97 |
| RB798 | *rrf-1(ok589)* | 120/140 | 118.33 | 2.88 | <0.0001 | |
| AGP273 | *rrf-1; tcer-1* | 127/148 | 102.84 | 2.38 | 0.0001 | 1 |
| **Trial 3** | | | | | | |
| N2 | Wildtype | 149/181 | 90.81 | 1.65 | | |
| CF1266 | *tcer-1(tm1452)* | 113/163 | 97.57 | 1.74 | 0.0354 | |
| NL3511 | *ppw-1(pk1425)* | 141/180 | 86.38 | 1.44 | 0.0624 | |
| AGP274 | *ppw-1; tcer-1* | 127/181 | 93.08 | 1.75 | 1 | 0.479 |
| RB798 | *rrf-1(ok589)* | 134/180 | 96.04 | 1.51 | 0.3229 | |
| AGP273 | *rrf-1;tcer-1* | 103/154 | 99.44 | 1.76 | 0.007 | 0.3859 |

**Table S3.** Impact of loss of small RNA biogenesis factors on worm survival on PA14.

| Strain | Genotype | n = obs/total | Mean (hrs) | SEM | Bonferroni *p* (vs. N2) |
|---|---|---|---|---|---|
| **Trial 1** | | | | | |
| N2 | Wildtype | 137/155 | 118.27 | 2.7 | |
| CF2166 | *tcer-1(tm1452)* | 119/170 | 183.11 | 4.35 | <0.001 |
| NL1810 | *mut-16(pk710)* | 127/145 | 135.95 | 2.95 | <0.001 |
| GR1946 | *mut-14(pk738) smut-1(tm1301)* | 139/159 | 169.7 | 4.52 | <0.001 |
| NL1820 | *mut-7(pk720)* | 129/149 | 138.18 | 4.44 | <0.001 |
| **Trial 2** | | | | | |
| N2 | Wildtype | 112/162 | 111.47 | 1.92 | |
| CF2166 | *tcer-1(tm1452)* | 120/201 | 151.35 | 4.2 | <0.001 |
| NL1810 | *mut-16(pk710)* | 110/141 | 116.67 | 3.14 | 0.6114 |
| GR1946 | *mut-14(pk738) smut-1(tm1301)* | 134/160 | 141.88 | 3.7 | <0.001 |
| NL1820 | *mut-7(pk720)* | 89/115 | 160.94 | 5.52 | <0.001 |
| WM30 | *rde-3(ne298)* | 124/153 | 101.24 | 1.64 | <0.001 |
| **Trial 3** | | | | | |
| N2 | Wildtype | 117/158 | 137.64 | 3.34 | |
| CF2166 | *tcer-1*(tm1452) | 94/167 | 166.53 | 4.48 | <0.001 |
| NL1810 | *mut-16(pk710)* | 80/122 | 134.79 | 3.51 | 1 |
| DCL565 | *rde-1(mkc36)* | 115/164 | 127.83 | 3.04 | 0.1937 |
| WM45 | *rde-1(ne300)* | 105/134 | 102.26 | 1.27 | <0.001 |
| WM27 | *rde-1(ne219)* | 105/135 | 115.25 | 2.01 | <0.001 |
| **Trial 4*** | | | | | |
| N2 | Wildtype | 99/138 | 58.56 | 1.53 | <0.001 |
| CF2166 | *tcer-1(tm1452)* | 52/136 | 77.52 | 2.93 | <0.001 |
| NL1810 | *mut-16(pk710)* | 121/135 | 64.21 | 1.39 | 0.0309 |
| GR1946 | *mut-14(pk738) smut-1(tm1301)* | 115/138 | 59.46 | 1.66 | 1 |
| **Trial 5** | | | | | |
| N2 | Wildtype | 108/130 | 74.9 | 1.34 | |
| SX2499 | *prde-1(mj207)* | 71/104 | 75.84 | 2 | 1 |
| CF2166 | *tcer-1 (tm1452)* | 91/120 | 103.91 | 2.45 | <0.001 |

* PA14 exposure at 20˚C except trial 4 which was done at 25˚C.
